# Growth-limiting drought stress induces time-of-day dependent transcriptome and physiological responses in hybrid poplar

**DOI:** 10.1101/2021.12.22.473933

**Authors:** Sean M. Robertson, Solihu Kayode Sakariyahu, Ayooluwa J. Bolaji, Mark F. Belmonte, Olivia Wilkins

## Abstract

Drought stress negatively impacts the health of long-lived trees. Understanding the genetic mechanisms that underpin response to drought stress is requisite for selecting or enhancing climate change resilience. We aimed to determine how hybrid poplars respond to prolonged and uniform exposure to drought; how responses to moderate and more severe growth-limiting drought stresses differed; and, how drought responses change throughout the day. We established hybrid poplar trees (*Populus* x ‘Okanese’) from unrooted stem cutting with abundant soil moisture for six weeks. We then withheld water to establish well-watered, moderate, and severe growth-limiting drought conditions. These conditions were maintained for three weeks during which growth was monitored. We then measured photosynthetic rates and transcriptomes of leaves that had developed during the drought treatments at two times of day. The moderate and severe drought treatments elicited distinct changes in growth and development, photosynthetic rates, and global transcriptome profiles. Notably, the time of day of sampling produced the strongest signal in the transcriptome data. The moderate drought treatment elicited global transcriptome changes that were intermediate to the severe and well-watered treatments in the early evening, but did not elicit a strong drought response in the morning, emphasizing the complex nature of drought regulation in long-lived trees.

**Highlight:** Poplar drought transcriptome is defined by the time of day of sampling and by the extent of water deficit.

## Introduction

Episodic drought can be catastrophic for forest trees and for the ecosystems they support (Assal *et al.*, 2016; Brodribb *et al.*, 2020). In response to drought, trees may slow their photosynthetic and stomatal conductance rates (Xiao *et al.*, 2009; Hamanishi *et al.*, 2012; Attia *et al.*, 2015), reduce their overall growth rates (Han *et al.*, 2019; Yi *et al.*, 2020), and experience increased rates of crown thinning and mortality (Worrall *et al.*, 2013). Because of their long lives, trees experience drought as a spatially and temporally heterogeneous stressor (Assal *et al.*, 2016). For example, drought responses are interwoven with underlying daily and seasonal changes in water demands (McLaughlin *et al.*, 2003), light availability (Aranda *et al.*, 2005), and changes in gene expression (Filichkin *et al.*, 2011). During periods of drought, trees develop morphologically distinct leaves with altered stomatal density (Hamanishi *et al.*, 2012; McKown *et al.*, 2014) and hormone profiles (Paul *et al.*, 2018) that reflect the water status of the plant during their development and which directly influence the ability of the tree to withstand a period of drought (Himes *et al.*, 2021). Additionally, because drought occurs over time, it is often coincident with other environmental stressors that may modulate drought response synergistically, additively, or antagonistically (Mittler, 2006; Plessis *et al.*, 2015).

Creating growth limiting soil water conditions in controlled environments like greenhouses and growth chambers that approximate drought conditions in nature is essential for studying physiologically relevant drought stress responses (Verelst *et al.*, 2013; Lovell *et al.*, 2016). Experiments that have directly compared drought stress responses in controlled and field environments have reported significant differences in physiological responses and gene expression profiles depending on the type of environments where plants were grown (Lovell *et al.*, 2016; Wilkins *et al.*, 2016). Some of these differences can be attributed to the degree or duration of the drought (Monroe *et al.*, 2021), or to the underlying complexity of the environment (Mittler, 2006; Plessis *et al.*, 2015). In nature, the onset of drought is often gradual, and plants adjust their physiology and development to minimize damage and optimize growth (Sohn *et al.*, 2013). In contrast, in controlled environments, trees are grown in relatively small volumes of soil and dry out relatively quickly (Wilkins *et al.*, 2009). As such, a water withholding treatment in controlled environments may be experienced as a shock rather than a drought and may elicit responses that diverge strongly from drought responses in the field (Lovell *et al.*, 2016). Moreover, long-lived trees in nature can use their large root volumes to exploit water from deeper soil horizons thereby extending the period of water sufficiency which may allow for the development of new tissues and organs during soil drying (Mackay *et al.*, 2020). Differences between responses in controlled and field environments are also critical considerations for genetic enhancement of stress tolerance emphasizing the need to study stress responses in conditions in which plants are grown (Zaidem *et al.*, 2019; Groen *et al.*, 2020).

Fluctuations in diel signals, including light, temperature, and atmospheric humidity have a substantial influence on plant water demand and drought responses. These environmental oscillations are linked to daily patterns of physiological and developmental processes, such as photosynthesis and respiration (Durand *et al.*, 2019) and leaf expansion (Pantin *et al.*, 2011). In addition to these physiological diel oscillations, the transcriptome also varies throughout the day, even while soil water conditions remain constant (Filichkin *et al.*, 2011). These predictable daily transcription patterns, together with oscillations in water demand, result to distinct drought transcriptomes at different times of day (Wilkins *et al.*, 2009; Hamanishi *et al.*, 2010; Dubois *et al.*, 2017). Thus, to capture the full spectrum of stress responses elicited in poplars, drought-responsive alterations in physiology and gene expression must be examined at multiple times of the day.

Trees of the genus *Populus* are well suited for studying molecular drought responses as they are propagated clonally, have a high-quality reference genome (Tuskan *et al.*, 2006), and vary with respect to their tolerance to drought (Barchet *et al.*, 2014; Attia *et al.*, 2015). Moreover, trees of this genus are foundational to many temperate ecosystems and are important as a short rotation fiber crop and for carbon sequestration throughout the Northern hemisphere (Fang *et al.*, 2007). Understanding the genetic mechanisms underpinning these responses and which create the adaptive variation in drought tolerance in forest trees may illuminate potential avenues for enhancing drought tolerance (Estravis-Barcala *et al.*, 2020). To gain broader insight into how poplar trees respond to drought stress, we designed an experiment to overcome some of the limitations of controlled environment drought experiments. We grew hybrid poplar trees (*Populus* x ‘Okanese’) (Schroeder *et al.*, 2013) in greenhouse conditions for 42 days under well-watered conditions. We then transferred the trees to one of three gravimetric soil water conditions for three weeks. After the trees had been acclimated to the new conditions, we measured a variety of plant growth and photosynthesis parameters, in addition to gene expression profiles in fully expanded leaves to characterize elements of the drought response. The aims of this work were to determine how hybrid poplar respond to prolonged and uniform exposure to water limiting conditions; to determine if the responses to moderate and more severe drought stresses were qualitatively or quantitatively different; and, to determine how the response to drought changes throughout the day.

## Materials and Methods

### Plant Growth Conditions

Unrooted stem cuttings (*Populus* x ‘Okanese’) were soaked in water for 24h at 4°C in the dark, trimmed to remove dead ends, and planted in 4-liter pots filled with Sungro Sunshine Professional Growing Mix (Sun Gro Horticulture, Agawam, Massachusetts). Trees were grown in a climate-controlled greenhouse with 22-24°C days and 15-17°C nights and a photoperiod of 15h. All trees were fertilized with Miracle-Gro All-Purpose Water-Soluble Plant Food 24-8-16 (MiracleGro, Marysville, Ohio) once before the water-withholding treatments were initiated.

### Sample Collections

To ensure that all measurements were made on fully expanded leaves that developed entirely during the treatment period, the uppermost leaf on each tree was tagged 42 days after planting. On this day, all trees were assigned to one of three treatment groups with gravimetric soil water contents of 30%, 50%, or 80% of field capacity. For the subsequent 21 days, pots were weighed daily and watered as needed to maintain them at their target soil water contents (Fig. S6). Photosynthetic measurements were made 21 days after water treatments were initiated, once between 10:00 and 12:00, and again between 14:00 and 16:00, using a Li-Cor LI-6400XT Infrared Gas Analyzer (LI-COR, Lincoln, Nebraska). Leaves for transcriptome analysis were harvested at 10:00 (late morning) and 16:00 (early evening), flash-frozen in liquid nitrogen, and stored at −80°C until further processing.

### Image analysis

The area of leaves that had developed and expanded during the treatments were measured using PlantCV (Gehan *et al.*, 2017). Leaf images were captured in a lightbox using an iPhone-X camera. The images were processed using the multi-plant workflow to estimate leaf area. First, the leaf images were masked from the background pixels to isolate the leaf pixels and then, the area was quantified in pixels using the shape_img function of PlantCV. Analysis of variance was carried out in R to determine statistical significance between the leaf area of the treatment groups.

### Transcriptome Library Preparation and Sequencing

Leaf tissue was ground to a fine powder under liquid nitrogen and total RNA was extracted using RNeasy Plant MiniKits (Qiagen, Toronto, Canada). RNA was sent to Genome Québec (Montréal, PQ, Canada) where NEBNext stranded sequencing libraries were prepared according to manufacturer’s instructions (New England Biolabs, Whitby, ON, Canada) and ingle end 100 base pair reads were sequenced on an Illumina HiSeq4000. A minimum of 23 million reads were generated for each library (Table S1). Raw and processed data are available for download as accession number GSE191155 on NCBI’s Gene Expression Omnibus (Edgar *et al.*, 2002).

### RNA-seq Data Analysis

Processing of the raw read files was accomplished using high-performance computing clusters provided by WestGrid (www.westgrid.ca) and Compute Canada (www.computecanada.ca). Read quality was assessed using FastQC (Andrews, 2010). Low-quality reads and adaptor sequences were removed with Trimmomatic (Bolger *et al.*, 2014). The reads were aligned to the *Populus trichocarpa* v4.1 reference genome (Tuskan *et al.*, 2006) (downloaded from the Phytozome database (Goodstein *et al.*, 2012)) ) using HISAT2 (Kim *et al.*, 2019) and converted to BAM format with SAMtools (Li *et al.*, 2009). Transcript abundance was quantified using featureCounts (Liao *et al.*, 2014). All downstream analyses were performed in R (R Core Team, 2021) using edgeR (Robinson *et al.*, 2010) and plotted using ggplot2 (Wickham, 2007). Genes were filtered at an average logCPM > −2 and < 9 to remove low gene counts and outliers, normalized using the trimmed mean of M-values method (Robinson *et al.*, 2010), and dispersion estimates were calculated using the estimateGLMRobustDisp function. Differentially expressed genes (DEGs) were identified as those with an absolute log_2_ fold change > 0 and a false discovery rate [FDR] < 0.05 using the Benjamini-Hochberg method (Benjamini and Hochberg, 1995). The code for the stacked bar plot (Fig. 5A) figure was modified from Bjornson et al., 2021(Bjornson *et al.*, 2021). Principal Component Analysis was performed using the plotPCA function in DESeq2 (Love *et al.*, 2014) and the Pearson correlation coefficient using cor.test function. All code for this project is available at https://github.com/Wilkins-Lab/Poplar2021.

### Clustering and GO term enrichment analysis

Transcript profiles were clustered using cutTreeDynamic from the package dynamicTreeCut (Langfelder *et al.*, 2008) using a minimum cluster size of 20 and a deepsplit value (sensitivity) of 1. Gene set enrichment analysis was performed using topGO (Alexa and Rahnenfuhrer, 2021), using Fisher’s exact test (*P* < 0.05). Gene Ontology terms (*Populus trichocarpa V3.0* GO annotation) were downloaded from agriGO v2.0 (Tian *et al.*, 2017). Only 21,638 genes (<50% of all genes) have annotated GO terms; as such, some functional categories may not be reported as enriched because of limitations on the GO annotation for Poplar.

## Results

### Growth and photosynthesis rates are reduced with declining soil water content

After 21 days growing in soil media with gravimetric soil water content (SWCg) of 20-30, 40-50, 70-80% of field capacity, trees in the 80% SWCg group were taller (Fig. 1A) and had developed more new leaves (Fig. 1B) than trees in the other two treatment groups. Notably, this was a graduated response, where trees in the 50% SWCg group were taller than trees in the 30% SWCg group in both measured aspects. The difference in total leaf area for trees between 50% and 80% SWCg groups (Fig.1D) is principally be the product of the number of leaves (Fig.1B) as the size of newly developed leaves in these treatment groups was indistinguishable (Fig. 1C). This is in contrast to trees in the 30% SWCg group, which developed fewer (Fig. 1B) and smaller (Fig. 1C) leaves than either of the other treatment groups. Additionally, the growth rate of the trees decreased upon water withholding, with the 50% and 80% SWCg groups stabilizing after ~14 days, while the 30% SWCg group growth rate continued to decline throughout the experiment (Fig S1A).

**Figure 1.**
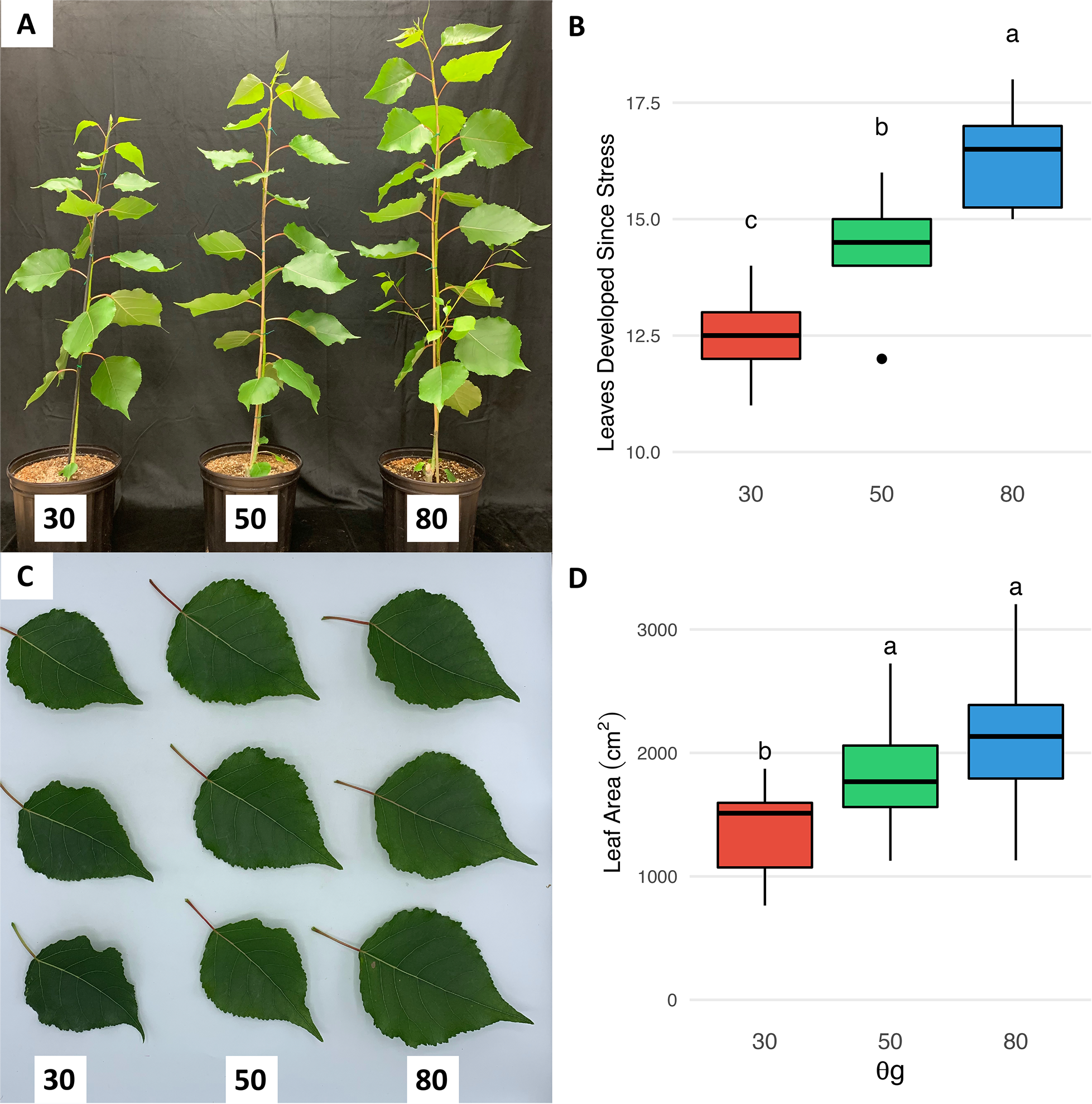
Moderate and severe growth-limiting drought induces a graduated reduction of poplar leaf and stem growth. Growth metrics were reduced in **A.** plant height (n=16), **B.** number of newly emerged leaves since the onset of stress (n=16), **C.** size of third leaf fully developed in stress (n=6), and **D.** area of third leaf fully developed in stress (n=6). Significance was determined by Tukey’s Honest Significant Difference test (P < 0.05).

To explore the physiological consequences of water limitation, we measured photosynthetic, transpiration, and stomatal conductance rates in the morning (10:00-12:00) and afternoon (14:00-16:00) of the 21^st^ day of the water withholding treatments (Fig. 2). All three of these parameters decreased gradually with decreasing soil water availability at both times of day (Fig. 2A-C); however, for a given treatment group, none of the parameters differed significantly between the morning and afternoon measurements (*P* < 0.05). Water use efficiency (WUE) was higher in trees in the 50% SWCg group than in trees in either of the other treatments (Fig. 2D). WUE was extremely variable for trees in the 30% SWCg treatment group, with both the highest and lowest calculated WUEs for any treatment group in the morning and afternoon. To explore the source of this variation, we plotted the relationship between the actual soil water content of each pot, which was below the upper limit of the target range at the time of the physiological measurement, and WUE (Fig. S1A). This analysis determined that WUE increased in the trees in the 30% SWCg group, as long as the overall weight of the wet soil media was greater than ~1000g; WUE rapidly declined if the water content was below this threshold. Because the photosynthetic parameters were measured before plants were re-watered to the upper limit of their soil water content, the photosynthesis measurements were made when plants may have been in even more water limiting conditions.

**Figure 2.**
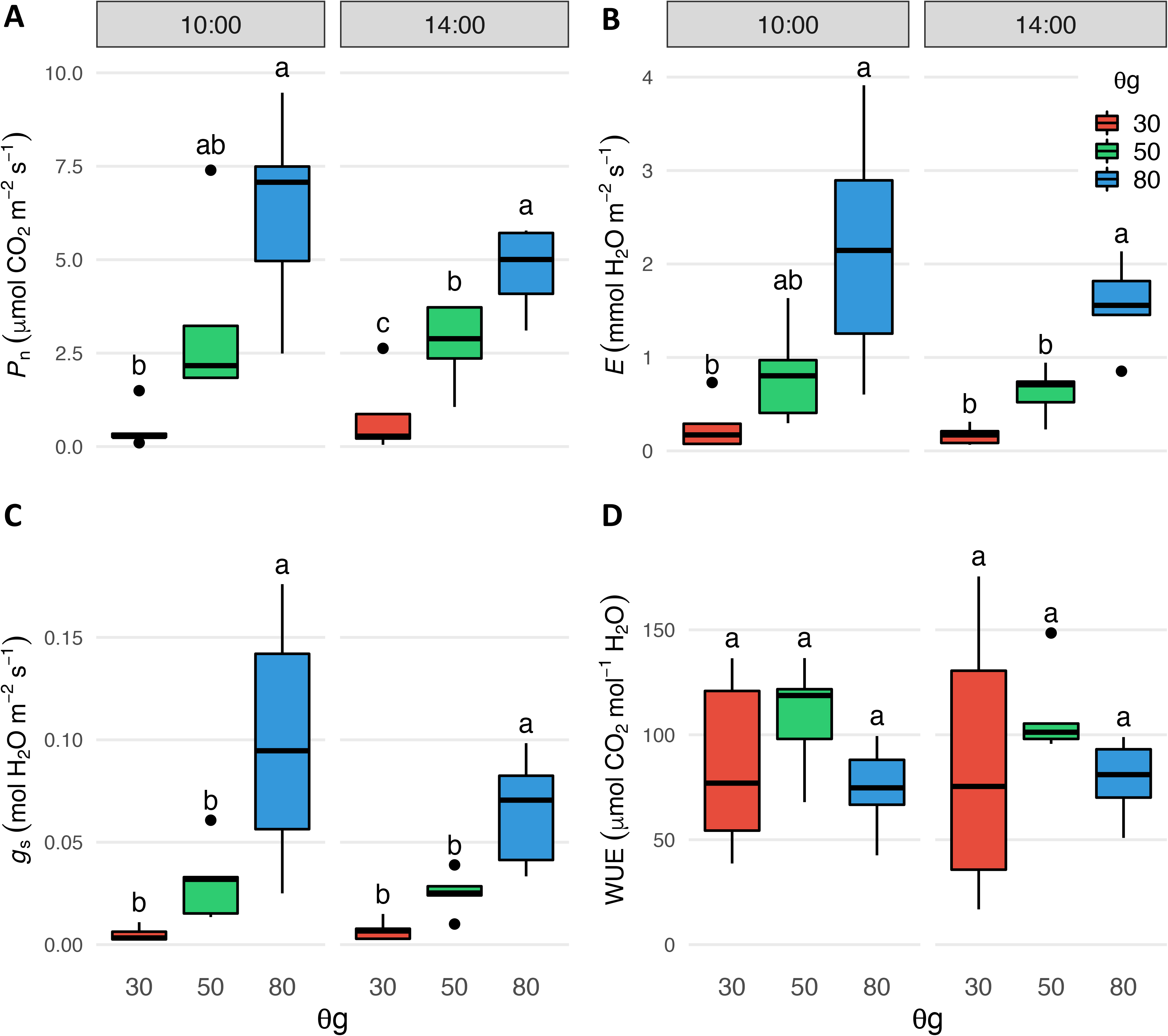
Photosynthesis, transpiration, and stomatal conductance rates decline with decreasing soil water availability. **A.** Net photosynthetic rate *P*_*n*_, **B.** transpiration rate *E*, **C.** stomatal conductance *g*_*s*_, and **D.** water-use efficiency (WUE) of poplar leaves (n = 5 leaves/treatment/timepoint) Measurements were taken between 10:00 - 12:00 and again between 14:00 - 16:00. Water-use efficiency was calculated as the ratio of *P*_*n*_ to *g*_*s*_. Significance was determined by Tukey’s Honest Significant Difference test (*P* < 0.05).

### Time of day is the strongest source of variation in the transcriptome

To understand molecular mechanisms underlying the long-term response to water deficit, we measured the transcriptomes of leaves harvested from hybrid poplar trees grown in three soil water content treatments at two times of day as described above. In total, more than 600 million 100bp single-end reads were generated across 18 samples (Table S1). Approximately 90% of reads passing the quality threshold (‘surviving reads’) mapped to unique loci in the *Populus trichocarpa* v4.1 reference genome (Tuskan *et al.*, 2006) (Table S1). To identify the main sources of variation in the transcriptome data, we conducted a principal component analysis (PCA) on normalized read counts. The first principal component (PC1) explained 94% of the variance in the transcriptome data (Fig. 3A). The samples were cleanly divided along this axis by the time of day at which they were collected, indicating that time of day was the greatest source of variation in the transcriptome data. The second component (PC2) explained 2% of transcriptome data variation (Fig. 3A). The samples are imperfectly stratified by the water treatment group along this axis. The transcriptomes of leaves from 30% SWCg were strongly separated from the transcriptomes of the other treatment groups in the late morning; this separation was less apparent in the transcriptome data from the early evening. Additionally, the third principal component accounted for 1% of the variation observed in our dataset (Fig S3).

**Figure 3.**
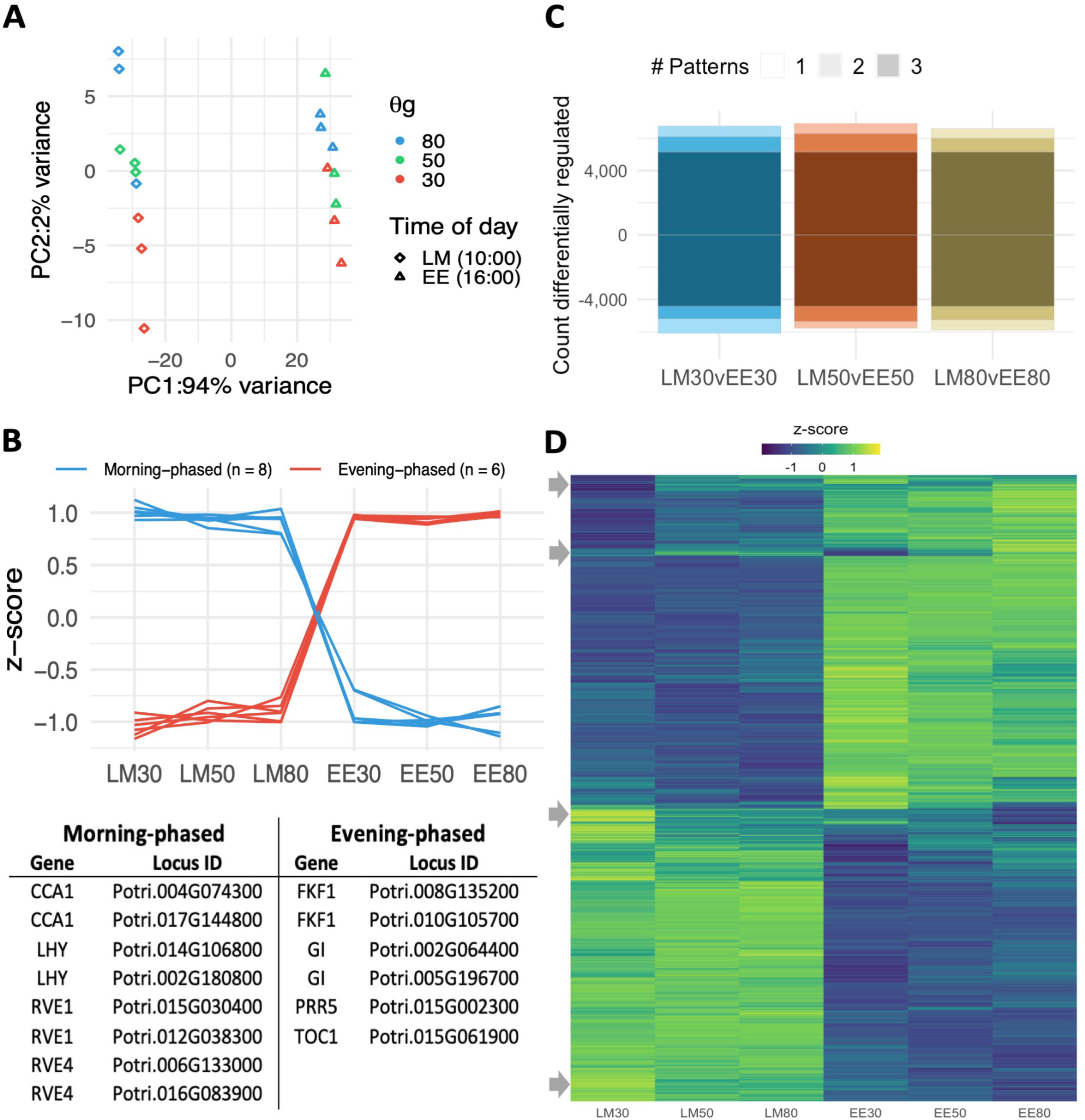
The time of day of sampling produced the strongest signal in the leaf transcriptome data. **A.** Principal component analysis of transcript profiles of all samples. Shape of data point indicates time of sampling; colour corresponds to soil water treatment. **B.** Expression profiles of selected clock genes across all treatments, gene name and locus of clock genes are listed below the expression profiles **C.** Number of differentially expressed genes between samples collected from plants in the same soil water treatment at different times of day (|log_2_fc| > 0, FDR < 0.05) **D.** Heatmap of the 13269 differentially expressed from C. The arrows identify clusters of genes whose expression profiles are influenced by both time of day of sampling and water status.

Given the strong time-of-day signal in the data set, we mapped the transcript abundance profiles of genes with well-characterized daily expression patterns to explore their expression in the present experimental conditions. We plotted the transcript abundance of 14 circadian clock genes, including eight morning-phased and six evening-phased genes (Fig. 3B). As expected, morning-phase genes displayed higher abundance in the late morning, and the evening-phase genes had consistently higher abundance in the early evening (Fig. 3B). For each water treatment, transcripts that were differentially expressed between the two sampling times (Fig. 3C; False discovery rate [FDR] - adjusted *P* < 0.05)) were identified. In total, 13,269 differentially expressed genes (DEGs) were identified of which approximately half were higher in the morning, and half were higher in the evening (Fig. 3C, Table S2). More than 9,500 genes were differentially expressed between the sampling times in all treatment groups, while only 3,833 genes were differentially expressed at a single level of drought stress (Fig. 3C, Table S2). While most genes that were differentially expressed between the morning and evening samples were unaffected by the soil water conditions, several small groups of transcripts were affected by both the time of sampling and by water availability (Fig. 3D).

### Differentially expressed genes at each time of day have distinct functions in the drought stress response

For each time of day, we identified differentially expressed genes (DEGs) between the drought treated (30% and 50% SWCg) and the control samples(80% SWCg, FDR < 0.05). In total, 971 DEGs were identified between the late morning samples, and 1,448 DEGs were found between the early evening (Fig. S3). Transcript profiles were then hierarchically clustered using the complete linkage method, resulting in 960 of the morning DEGs and 1423 of the evening DEGs assigned to clusters. Clusters that included more than 20 genes and had highly correlated (pairwise correlation > 0.99) transcript profiles were further characterized (Figure 4A, 5B; genes in each cluster are included in Fig. S3). Amongst these clusters, four principal expression patterns were observed (Figure 4C-D): transcripts that had increased abundance only at the lowest level (30% SWCg) of water availability (patterns I: 226 genes, and VI: 27 genes), transcripts whose abundance increased gradually with decreasing soil water availability (patterns II: 227 genes, and VII: 560 genes), transcripts that had decreased abundance only at the lowest level (30% SWCg) of water availability (patterns III: 389 genes, and VIII: 40 genes), and transcripts whose abundance decreased gradually with decreasing water availability (patterns IV: 118 genes, and IX: 631 genes). In the early evening DEGs, however, two additional distinct patterns are seen: upregulated and downregulated in low and moderate water availability (pattern V: 66 genes, and X: 99 genes). These two patterns account for 12% (165/1423) of the clustered early evening genes. Additionally, in the early evening, 84% (1,191/1,423) of clustered DEGs had a graded response to water availability, wherein transcripts in the 50% SWCg accumulated to a moderate level between the 30% and 80% SWCg values (pattern VI: 560 genes; pattern IX: 631 genes). In contrast, in the late morning, 64% (615/960) of the clustered genes were increased (pattern I) or decreased (pattern III) only in the 30% SWCg treatments.

**Figure 4.**
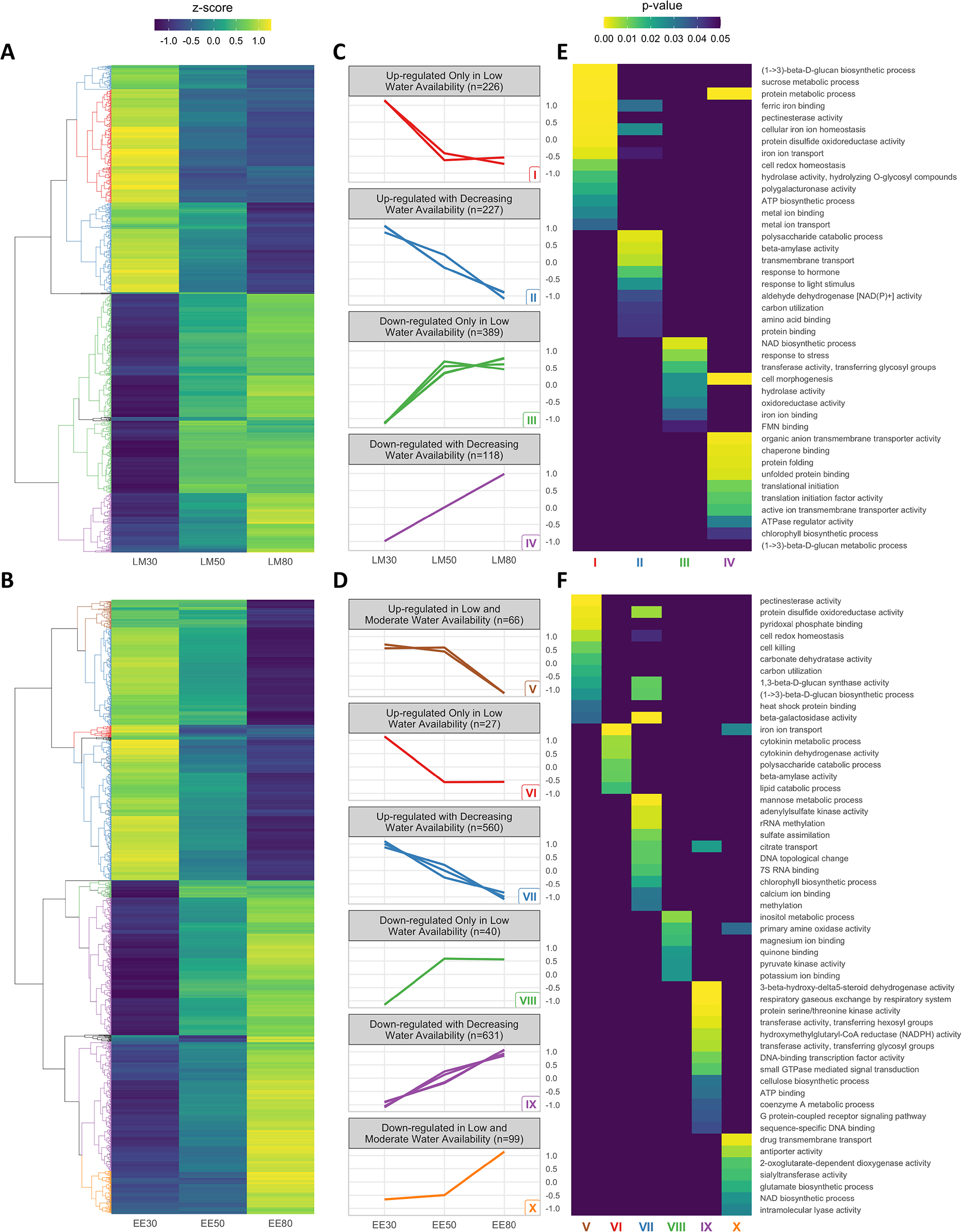
The extent of water deficit and time of day induce distinct patterns of gene expression with diverse molecular functions. **A-B.** Differentially expressed genes each of the drought stress treatments and the control samples (FDR < 0.05) were hierarchically clustered according to scaled expression profiles. A. Heatmap of DEGs in the late morning samples (LM, n=981) and B. Heatmap of DEGs in the early evening samples (EE, n=1464). Dendrograms are coloured to highlight the clusters further characterized in C and D. **C-D.** Scaled expression profiles of clusters within each of the expression patterns; each line is the mean expression of all transcripts in one cluster identified in A or B. **E-F.** Selected enriched Gene Ontology terms within each of the expression patterns identified in C and D (*P* < 0.05).

We next performed a gene set enrichment analysis of each of the clustered gene patterns. The enriched Gene Ontology (GO) terms revealed distinct functional responses within the gene expression patterns of DEGs (Fig. 5E-F; Supp. Fig. S4). Notably, we found that functionally different gene sets were enriched for the same expression patterns observed in the morning and evening. For example, genes that were downregulated with decreasing water availability were enriched for translational initiation and post-translational modification in the morning, but cell signalling and DNA binding in the evening. Moreover, we show that common GO terms were enriched in different expression profiles in the morning and evening samples. For instance, functions related to cell wall modification were expressed only in low water availability in the morning, but in both low and moderate water availability in the evening (Fig. 5).

**Figure 5.**
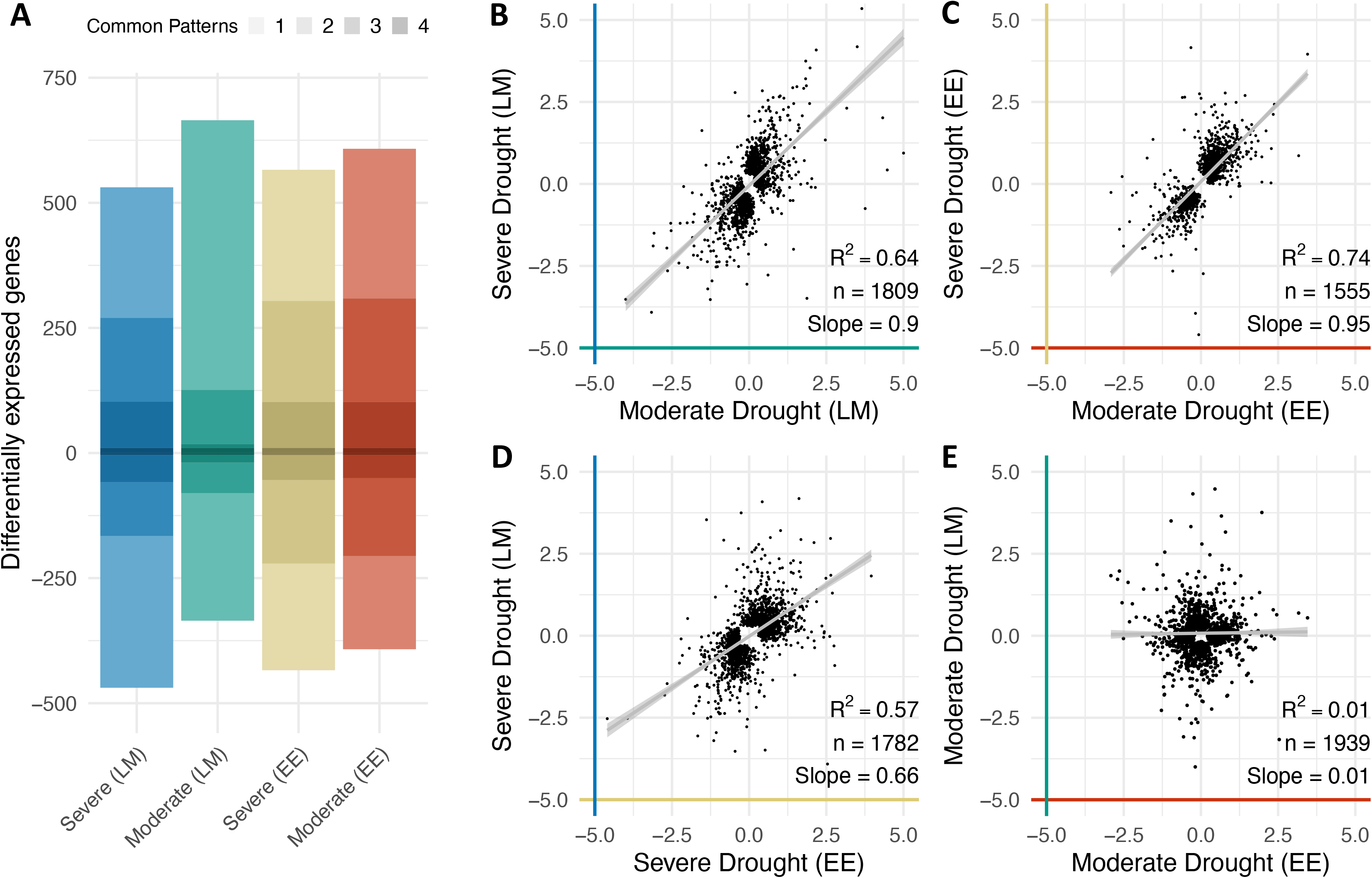
The extent of water deficit and time of day interact to induce distinct transcriptome-wide profiles, indicating condition dependent quantitative shifts in gene expression. **A**. The top 1000 up and downregulated genes (ranked by FDR) in severe and moderate drought at each time of day. Intensity of the bar color indicates the number of conditions in which a given gene is differentially expressed. **B-E** Pearson correlation of expression changes between pairs of treatments: severe and moderate water stress at late morning (B), and early evening (C), and the late morning and early evening for moderate (D) and severe drought (E) Linear regression line shown in gray, with Pearson correlation coefficient (R2), number of genes, and slope of the regression line shown on each plot. The colors of the axes match the colours used in panel A.

### Global analysis of time-of-day transcriptome identifies poplar response to water stress

To determine if any genes were universally responsive to the water stress treatment in our experiments, we selected the top 1000 DEGs (FDR rank-ordered) for each water stress treatment (30% and 50% SWCg) compared to the control (80% SWCg) at each time of day (Table S3). This analysis found just over half of the possible 4000 genes that could be generated by this analysis (2,362 genes) were differentially expressed in only one contrast (Fig. 5A). The remaining genes were differentially expressed in two (565), three (152 genes), or all four (13 genes) contrasts (Fig. 5A; Table S4). The list of differentially expressed genes identified in the late morning 50% vs. 80% SWCg contrast was dissimilar to the other three DEG lists; more than 80% of DEGs (812/1000) on this list were unique to this treatment. Of the 565 DEGs that were common to two patterns, 435 (77%) were common to different treatments at the same time of day.

To explore the quantitative relationships between the milder and more severe water limiting treatments, we calculated Spearman’s rank correlation coefficient for the top 1000 DEGs for each treatment as described above. We found a weaker correlation between the expression of the genes in response to milder (50% SWCg) and more severe water deficits (30% SWCg) in samples collected in the morning (rho=0.65, Fig. 5B) compared to samples collected in the evening (rho = 0.81, Fig. 5C), although the slope of these relationships was similar. We also determined a moderate correlation between the expression of DEGs in the more severe water limiting conditions (30% SWCg) between the morning and evening (rho=0.64, Fig. 5D), while no relationship existed between the expression of DEGs in the milder water limiting conditions (50% SWCg) across sampling time (rho=0.01, Fig. 5E).

## Discussion

### Plants can adapt to water limiting conditions during prolonged exposure

In the current study, we show that reduced water availability slows stem growth in hybrid poplar and that the rate of growth stabilizes after 2-3 weeks in the tested conditions. This finding is consistent with a poplar drought study that found growth rates stabilized following a 15-day water deficit stress. Previous studies on drought stress in poplars have also shown that radial stem growth continued to decline over 11 days of water withholding (Bloemen *et al.*, 2016) and that the overall growth rate did not stabilize until 8-15 days after the water deficit was initiated (Bogeat-Triboulot *et al.*, 2019). In our study, growth rate stabilization was concomitant with robust decreases in photosynthetic and stomatal conductance rates in plants grown with limited water availability. Studies in poplar (Attia *et al.*, 2015), Kentucky bluegrass (*Poa pratensis*)(Saud *et al.*, 2016), and *Arabidopsis thaliana* (Bechtold *et al.*, 2016) that have measured photosynthesis and gas exchange parameters throughout longer-term soil drying experiments have shown that significant changes in these parameters occur only after plants have acclimated to their new conditions. In this study, we showed that leaves that developed and matured entirely during stress conditions were smaller and fewer than leaves that developed without water limitation. Detailed studies of the effects of drought on poplar leaf development have demonstrated that drought leads to variations in morphological and anatomical features including thicker leaf tissues, increased stomatal density (Regier *et al.*, 2009), and reduced leaf size (Hamanishi *et al.*, 2012; Viger *et al.*, 2016), in addition to biochemical changes, such as total non-structural carbohydrate concentrations (Regier *et al.*, 2009). Such differences in leaves give rise to differences in photosynthetic parameters (Cao *et al.*, 2012) and responses to water availability (McKown *et al.*, 2014). Temporal and spatial variation in the physiological and morphological features of leaves during drought, suggest that molecular measurements made during the initial period of stress may reflect transitional states rather than adaptive responses to stressful environmental conditions.

Despite the clear effects that water limitation had on the growth and gas exchange parameters in the current study, global RNA sequencing revealed the number of genes that were differentially expressed and the magnitude of gene activity were lower than had been reported in previous transcriptome drought studies in poplar (Wilkins *et al.*, 2009; Hamanishi *et al.*, 2010; Raj *et al.*, 2011). We hypothesized that this difference may exist because, unlike in the previous studies, the current study measured the transcriptomes of leaves that developed in water limiting conditions, where the trees had time to acclimate to the soil water treatments before sampling. However, the results of our transcriptomic analysis are consistent with a study in switchgrass (Lovell *et al.*, 2016) that identified fewer DEGs after long-term drought than in short-term dry down experiments. These disparities may be a result of differential gene expression between short-term and long-term drought responses. This explanation is consistent with the concurrent stabilization of growth and changes in photosynthetic parameters that we observed in the current study. Although trees may experience abrupt changes in water availability in the field, longer-term water limitations due to erratic precipitation patterns are more ecologically relevant to tree growth and productivity (Brodribb *et al.*, 2020).

### Time of day and the extent of drought stress affect photosynthesis and global changes in gene expression

In the current study, as in other studies of drought response in poplar, distinct patterns of gene expression were invoked depending on the time of day of observation (Wilkins *et al.*, 2009; Hamanishi *et al.*, 2010), in addition to the extent of water deficit experienced by the trees (Yan *et al.*, 2012; Cao *et al.*, 2014). By far, the largest source of variation in gene expression in the present study was the time of day of sampling. This was anticipated since an estimated 30-50% of *Arabidopsis* genes (Bla◻sing *et al.*, 2005; Covington *et al.*, 2008), 74% of *Brassica rapa* genes (Greenham *et al.*, 2020), and 60% of poplar genes (Wilkins *et al.*, 2009; Filichkin *et al.*, 2011) have predictable and reproducible daily changes in transcript abundance in non-stress conditions. Responses to any additional perturbations, such as environmental stressors, must be invoked on this underlying, dynamic transcriptional landscape (Greenham and McClung, 2015; Bonnot and Nagel, 2021). Studies in poplar (Wilkins *et al.*, 2009; Hamanishi *et al.*, 2010; Raj *et al.*, 2011), *Arabidopsis* (Wilkins *et al.*, 2010; Dubois *et al.*, 2017; Blair *et al.*, 2019), and rice (*Oryza sativa*) (Wilkins *et al.*, 2016; Grinevich *et al.*, 2019; Desai *et al.*, 2021) have shown that transcriptional responses to stress are different at different times of day. This was consistent with our findings here that the majority of drought responsive genes had distinct expression profiles in the morning and early evening. Our observations may be attributable to underlying diurnal variation in expression patterns. For example, global changes in gene expression in response to water limitation or a given level of water availability are perceived as a stressor only at some times of day (McLaughlin *et al.*, 2003).

Here we show that different functional classes of drought responsive genes were enriched at different times of day and with different degrees of water limitation. For example, genes involved in the regulation of iron transport were induced by water limitation in the morning only. The transport of iron from the vasculature to the chloroplasts is regulated by the circadian clock (Salomé *et al.*, 2013), and is affected by water availability (Pan *et al.*, 2015). We also showed that genes involved in protein folding and translation initiation were reduced in water-limited samples in the morning, but were unchanged in response to drought later in the day. This finding may be consistent with the growing literature showing that stress responses may be regulated in part by translation (Reynoso *et al.*, 2019; Bonnot and Nagel, 2021) and post-translational modifications (Rodriguez *et al.*, 2021). Together, these findings highlight the importance of including multiple timepoints in experimental design for capturing the full suite of stress response mechanisms.

In contrast to the strong time of day effect on gene expression profiles, we determined that water use efficiency (WUE) was most strongly affected by the extent of water limitation. We showed that WUE increased as water availability decreased in moderate moisture limiting conditions; however, in more severe water limiting conditions, WUE was highly variable and included high and low WUE values. As previously described in maize (Blankenagel *et al.*, 2018), in the present study there appeared to be a water content threshold below which WUE declined precipitously. This observation may be related to exponential decreases in soil water potential at low SWCg (Dowd *et al.*, 2019), whereby a small change in water volume leads to a large change in water availability. Moreover, physiological and growth traits, including specific leaf area and leaf carbon content, are affected non-linearly by soil water availability (Monroe *et al.*, 2021) highlighting the importance of studying these responses under a limited number of discrete soil moisture conditions. To fully characterize plant responses to drought stress, it will be necessary to incorporate intermediate water limiting conditions to understand how the degree of water stress impacts plant status, rather than an “all-or-nothing” approach.

### Response to drought stress relies on qualitative and quantitative changes in the transcriptome

Response to drought stress in poplar is not limited to a subset of high-amplitude drought-responsive genes; rather it is a transcriptome-wide adjustment that is affected by the time of day and by the extent of the stress applied. In this study, we show that drought-induced transcriptome profiles across different water levels and time of day lead to qualitative and quantitative changes, as seen in previous studies (Lovell *et al.*, 2016). Qualitative changes refer to the specific genes invoked, as the poplars expressed different genes depending on their water status and time of day, while quantitative changes are represented by the degree of change observed, as the water status induced variable proportions of genes. We find that genes differentially expressed in more severe water limiting conditions were also differentially expressed, though not significantly so, in the moderate water limiting conditions at both times of day, and that this relationship was stronger in the early evening than in the late morning. Notably, low water availability in the morning and evening elicited similar degrees of differential expression across a wide panel of genes, but there was no correlation between differentially expressed genes in moderate water availability at the two times of day.

These findings demonstrate that water availability, diel signals, and the endogenous circadian clock interact to determine what conditions will be perceived as stress and which biological mechanisms will be invoked in response to environmental perturbations. Previous studies in *Arabidopsis* (Legnaioli *et al.*, 2009) and soybean (Marcolino-Gomes *et al.*, 2014) have shown the interconnectivity of the circadian clock and the drought response, likely due to the reciprocal regulation of abscisic acid with clock components (Legnaioli *et al.*, 2009; Pokhilko *et al.*, 2013). Widespread changes in the timing, magnitude, and diversity of gene regulation are necessary for plants to adequately respond to stress (Greenham and McClung, 2015) and so experimental studies must take this complexity into account.

## Supplementary data

Fig. S1. The growth rate of poplars stabilizes at 14 days after initiation of water deficit, except in 30% SWCg conditions.

Fig. S2. Water use efficiency (WUE) of 3 SWCg conditions on the last day of water deficit show variability in the 30% SWCg group.

Fig. S3. The second and third principal components (PC) from the PCA performed in Fig. 3A.

Fig. S4. Hierarchical clustering and scaled gene expression profiles of DEGs in the (A) late morning (LM) and (B) early evening (EE). Expression profiles are ordered according to the colored bars from left to right. The mean expression of the cluster is represented by the red line.

Fig. S5. Full list of Gene Ontology (GO) terms for the (A) late morning and (B) early evening gene expression profiles. See Fig. 5 for list of patterns (I-X).

Fig. S6. Pot weight of poplars after initiation of water deficit. Rewatering occurred after poplars were below their target weight.

Table S1. Processing of raw RNA-sequencing reads. Surviving and dropped reads are from Trimmomatic, the overall alignment rate is from HISAT2, and overall gene assignment rate is from featureCounts.

Table S2. Scaled expression of all differentially expressed genes between the late morning (LM) and early evening (EE).

Table S3. Top 1000 differentially expressed genes (FDR-ranked) from each water stress treatment compared to control.

Table S4. Log_2_ fold-changes of shared top 1000 differentially expressed genes (FDR-ranked) between comparisons of time-of-day and extent of water stress.

## Abbreviations

SWCg: Gravimetric soil water content
WUE: Water use efficiency
DEG: Differentially expressed genes
GO: Gene Ontology
FDR: False discovery rate
*P*_n_: Net photosynthesis
*g*_s_: Stomatal conductance
*E*: Transpiration rate

## Acknowledgements

We wish to thank Dr. R. Soolanakayahally and C. Stefner of the Agriculture and Agri-Canada Indian Head Research Farm for providing the biological materials; Dr. S. Renault for access and training on the LI-6400; and R. Chen, S. Campbell, G. Pagcaliwagan, J. Lee, M. Kang, J. Sangiovanni for assistance growing plants. This research was supported by grants from the Fonds de recherche du Québec - Nature et technologies and the Natural Sciences and Engineering Research Council of Canada awarded to OW.

## Author Contributions

All authors contributed to the project conceptualization and methodology. Funding was acquired by OW, while supervision, project administration, and resources were provided by MFB and OW. Data curation, formal analysis, and software programming were performed by SMR, SKS, and AB. SR conducted the investigation, and SR and KS carried out validation, visualization, and drafting the manuscript. Review and editing of the manuscript were completed by MFB and OW.

## Data Availability

Transcriptome data are available for download from the Gene Expression Omnibus as accession GSE191155 (https://www.ncbi.nlm.nih.gov/geo/query/acc.cgi?acc=GSE191155). All code used in data analysis and figure generation available from https://github.com/Wilkins-Lab/Poplar2021.

